# Structure of the nucleotide exchange factor eIF2B reveals mechanism of memory-enhancing molecule

**DOI:** 10.1101/222257

**Authors:** Jordan C. Tsai, Lakshmi E. Miller-Vedam, Aditya A. Anand, Priyadarshini Jaishankar, Henry C. Nguyen, Adam R. Renslo, Adam Frost, Peter Walter

**Affiliations:** Howard Hughes Medical Institute; Department of Biochemistry and Biophysics, University of California at San Francisco, San Francisco, CA, USA; Department of Pharmaceutical Chemistry, and Small Molecule Discovery Center, University of California at San Francisco, San Francisco, CA, USA; Chan Zuckerberg Biohub, San Francisco, CA, USA

## Abstract

Regulation by the integrated stress response (ISR) converges on the phosphorylation of translation initiation factor eIF2 in response to a variety of stresses. Phosphorylation converts eIF2 from substrate to competitive inhibitor of its dedicated guanine nucleotide exchange factor, eIF2B, inhibiting translation. ISRIB, a drug-like eIF2B activator, reverses the effects of eIF2 phosphorylation and, remarkably, in rodents enhances cognition and corrects cognitive deficits after brain injury. To determine its mechanism of action, we solved an atomic-resolution structure of ISRIB bound in a deep cleft within decameric human eIF2B by electron cryo-microscopy. Structural and biochemical analyses revealed that formation of fully active, decameric eIF2B holoenzyme depended on the assembly of two identical tetrameric subcomplexes, and that ISRIB promoted this step by cross-bridging a central symmetry interface. Regulation of eIF2B assembly emerges as a rheostat for eIF2B activity that tunes translation during the ISR and that can be further modulated by ISRIB.

## Main Text

Protein quality control is essential to the maintenance of cellular and organismal health. To prevent the production of deleterious proteins, such as those from invading viruses or those produced in misfolding-prone environments, cells regulate protein synthesis. By arresting or accelerating the cardinal decision of translation initiation, cells effect proteome-wide changes that drive organismal functions, such as development, memory, and immunity (*1–3*).

A key enzyme in the regulation of protein synthesis is eukaryotic translation initiation factor 2B (eIF2B), a dedicated guanine nucleotide exchange factor (GEF) for translation initiation factor 2 (eIF2). eIF2B is composed of five subunits (α,β,γ,δ,ε) that assemble into a decamer composed of two copies of each subunit (*4–8*). The eIF2Bε subunit contains the enzyme’s catalytic center and associates closely with elF2Bγ (*9*). Two copies each of the structurally homologous elF2Bα, β, and δ subunits form the regulatory core that modulates eIF2B’s catalytic activity (*10–12*). eIF2B’s substrate, eIF2 is composed of three subunits (α,β,γ) and binds methionine initiator tRNA and GTP to form the ternary complex required to initiate translation on AUG start codons. eIF2’s γ subunit contains the GTP-binding pocket (as reviewed in (*13, 14*)).

In response to various inputs, many of which are cell stresses, phosphorylation of elF2α at serine 51 converts eIF2 from a substrate for nucleotide exchange to a competitive inhibitor of eIF2B. Phosphorylated eIF2 binds to eIF2B with enhanced affinity, effectively sequestering the limiting eIF2B complex from engaging unphosphorylated eIF2 for nucleotide exchange (*10–12*). Such inhibition leads to an attenuation of general translation and, paradoxically, the selective translation of stress-responsive mRNAs that contain small upstream open reading frames. This latter set includes mRNAs that encode transcriptional activators such as ATF4 (*15, 16*). In this way eIF2 phosphorylation elicits an intricate gene expression program. This pathway was termed the “integrated stress response”, following the discovery of several kinases that all phosphorylate eIF2α at serine 51 to integrate different physiological signals such as the accumulation of misfolded proteins in the lumen of the endoplasmic reticulum, the accumulation of doublestranded RNA indicative of viral infection, the cell’s redox status, and nutrient availability (*17*).

We previously identified an ISR inhibitor (ISRIB) that reverses the effects of eIF2α phosphorylation, restoring translation in stressed cells and blocking translation of ISR-activated mRNAs, such as ATF4 (*18, 19*). When administered systemically to wild-type rodents, ISRIB enhances cognition, leading to significant improvements in spatial and fear-associated learning (18). This remarkable effect relies on translation-dependent remodeling of neuronal synapses (*20*). eIF2 phosphorylation correlates with diverse neurodegenerative diseases and cancers, as well as normal aging (*21–24*). Additionally, a number of mutations that impair eIF2B activity lead to a neurodegenerative disorder of childhood known as vanishing white matter disease (VWMD) that is marked by cerebellar ataxia, spasticity, hypersensitivity to head trauma and infection, coma and premature death (25). As a well-characterized small molecule with rapid cross-blood-brain barrier equilibration, reasonable bioavailability, and good tolerability in rodent efficacy models, ISRIB and related analogs offer great potential for treating VWMD and a range of other devastating diseases that are currently bereft of therapeutic options (*18, 26*). Indeed in rodents, ISRIB entirely reverses cognitive deficits associated with traumatic brain injuries (*27*) and protects against neurodegeneration (*26*).

Previous work identified eIF2B as the molecular target of ISRIB (*28, 29*). ISRIB enhances eIF2B GEF activity three-fold, stabilizes a decameric form of the enzyme when analyzed in high salt conditions, and increases thermostability of eIF2Bδ (*28*). Mutations that render cells insensitive to ISRIB cluster in the N-terminal region of the eIF2Bδ subunit (*29*), and when projected onto the crystal structure of *S. pombe* eIF2B, two of the mutated residues map to its symmetric interface (*8*). These data hinted that ISRIB may activate eIF2B by binding near adjacent δ subunits to exert its blunting effects on the ISR. Here we report mechanistic and structural insights into ISRIB’s mechanism of action.

### ISRIB stabilizes decameric eIF2B, accelerating GEF activity

To investigate the mechanism by which ISRIB enhances the GEF activity of eIF2B, we engineered a recombinant *E. coli* expression system for co-expression of all five subunits of human eIF2B (Fig 1A). eIF2B purified as a monodisperse complex that sedimented at 13.6S, corresponding to the size of a decamer containing two copies of each subunit (Fig. 1B - AUC, Fig. S1A).

**Fig. 1:**
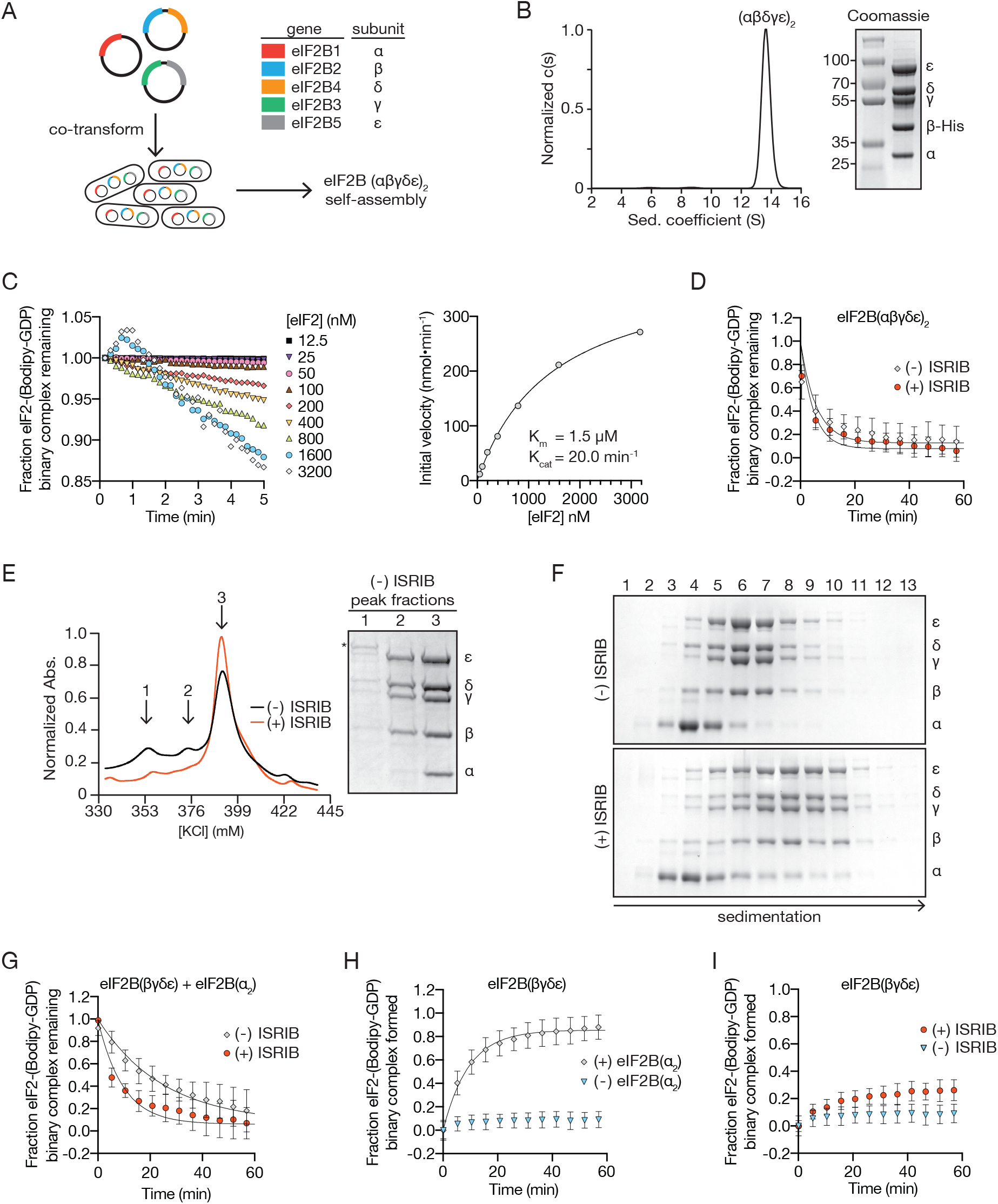
ISRIB stabilizes decameric eIF2B, accelerating GEF activity. (A) Schematic diagram for three plasmid expression of all five eIF2B genes in *E. coli*. (B) Characterization of eIF2B(αβγδε)_2_ by sedimentation velocity analytical ultracentrifugation and SDS-PAGE followed by Coomassie blue staining. (C) Initial rate of nucleotide exchange (right panel) plotted as a function of substrate concentration. Note that at high eIF2 concentration we reproducibly observed a transient increase in fluorescence that peaked at the 1 min time point (left panel). Such increase was reported previously (*29*) and remains unexplained. (D) GEF activity of eIF2B(αβγδε)_2_ as measured by unloading of fluorescent GDP from eIF2 in the presence and absence of ISRIB. (E) Absorbance 280 nm trace from an anion exchange column used in the purification of eIF2B in the presence (red) and absence (black) of ISRIB. Peak fractions from the (−) ISRIB purification were analyzed by SDS-PAGE and Coomassie-stained. eIF2B subunits are labeled (α-ε) and an asterisk denotes the presence of a contaminating protein that contributes to peak 1. (F) Stability of eIF2B(αβγδε)_2_ was assessed by sedimentation velocity on a 5–20% sucrose gradient in a 400 mM salt buffer. eIF2B(βγδε) and eIF2B(α_2_) were combined with and without 500 nM ISRIB. Fractions were analyzed by SDS-PAGE and Coomassie-stained. (G) GEF activity of eIF2B assembled from purified eIF2B(βγδε) and eIF2B(α_2_) in the presence and absence of ISRIB. (H) GEF activity of eIF2B(βγδε) in the presence and absence of eIF2B(α_2_). (I) GEF activity of eIF2B(βγδε) in the presence and absence of ISRIB.

We adapted a fluorescent GDP exchange assay (*29*), to assess the enzymatic activity of recombinant eIF2B. We purified the substrate, non-phosphorylated human eIF2, from a *S. cerevisiae* expression system genetically edited to lack the only yeast eIF2 kinase (*gcn2Δ*) (*30*) (Fig. S2A, S2B). First, in a ‘GDP loading assay’ we added fluorescent Bodipy-GDP to GDP-bound eIF2. We observed an eIF2B concentration-dependent increase in fluorescence corresponding to the dislodging of bound GDP and subsequent binding of Bodipy-GDP to eIF2 (Fig. S2C, Fig. S2D). Second, in a ‘GDP unloading assay’, we chased with a 1000-fold excess of unlabeled GDP and measured a decrease in fluorescence corresponding to the eIF2B-catalyzed dissociation of Bodipy-GDP from eIF2 (Fig. S2E). GEF activities were fit to a single-exponential (Fig. S2F) for calculating the reported k_obs_ values. Titrating substrate concentration to saturating levels in GDP unloading assays yielded K_m_ and k_cat_ values similar to those of eIF2B previously purified from mammalian cells (Fig. 1C) (*31*).

To investigate how ISRIB activates eIF2B, we fixed eIF2B and eIF2 in a multi-turnover regime at concentrations of 10 nM and 1 μM, respectively. Under these conditions, the eIF2 is subsaturating given its K_m_ of 1.5 μM (Fig. 1C). Previously, we reported a three-fold stimulation of nucleotide exchange by ISRIB under similar conditions (*28*). Surprisingly, ISRIB did not significantly activate the recombinant eIF2B decamer (Fig. 1D, (− ISRIB): k_obs_ = 0.17 +/− 0.006 min^−1^ and (+ ISRIB): k_obs_ = 0.21 +/− 0.005 min^−1^).

We previously showed that ISRIB stabilizes eIF2B decamers in lysates of HEK293T cells (28), suggesting a role during assembly of the active complex. To test this notion and its implications for ISRIB’s mechanism of action, we purified eIF2B in the presence or absence of ISRIB. Under both conditions we obtained the fully assembled decamer (Fig. 1E, peak 3); however, in the absence of ISRIB we also obtained a partially assembled complex lacking the a subunit that eluted from the anion exchange column at a lower ionic strength (Fig. 1E, peak 2). These data suggest that ISRIB enhances the stability of the decamer. To test this idea, we expressed eIF2B(βγδε) and eIF2Bα separately (Fig. S1B, Fig. S1C). Surprisingly, eIF2B(βγδε) purified as a heterotetramer, as determined by analytical ultracentrifugation (Fig. S1D), while eIF2Bα purified as a homodimer as previously observed (Fig. S1E) (*6*). We then combined eIF2B(βγδε) and eIF2B(α_2_) under stringent conditions of elevated ionic strength (400 mM) to assess ISRIB’s contribution to the stability of the decameric complex. When analyzed by velocity sedimentation in the absence of ISRIB, eIF2B(βγδε) sedimented as a tetramer (peak fractions 6–7) whereas eIF2B(α_2_) peaked in fraction 4 (Fig. 1F, upper panel). By contrast, in the presence of ISRIB, eIF2B(βγδε) and eIF2B(α_2_) sedimented together as a higher molecular weight complex deeper in the gradient (peak fractions 7–9) (Fig. 1F, lower panel). As we discuss below in Figures 3 and 4, the stabilized decamer peaks in fraction 10 of the gradient, indicating that under these conditions, the decamer partially dissociates during sedimentation. We surmise that dissociation during centrifugation led to the broad sedimentation profiles observed. Together these data show that ISRIB enhanced the stability of decameric eIF2B.

To understand the interplay between ISRIB binding, eIF2B(α_2_) incorporation into the decamer, and GEF activity, we mixed independently purified eIF2B(α_2_) and eIF2B(βγδε) subcomplexes and assayed the combination for GDP unloading. When assayed under these conditions, the specific activity was four-fold reduced when compared to the fully assembled decamer (compare Fig. 1D and 1G, k_obs_ = 0.17 +/− 0.006 min^−1^ and 0.04 +/− 0.009 min^−1^). Importantly, the addition of ISRIB restored GEF activity three-fold toward the level of fully assembled decamer (k_obs_ = 0.11 +/− 0.002 min^−1^) (Fig. 1G), suggesting that ISRIB’s activity reflects enhanced decamer stability.

Using the GDP loading assay, we found that eIF2B activity was reduced profoundly (k_obs_ = 0.01 +/− 0.007 min^−1^) in the absence of eIF2B(α_2_) (Fig. 1H), as previously reported (*32, 33*). Interestingly, ISRIB still activated eIF2B(βγδε) (Fig. 1I, k_obs_ = 0.04 +/− 0.003 min^−1^), indicating that ISRIB can enhance GEF activity independent of eIF2B(α_2_) incorporation into the holoenzyme. To reconcile these unexpected findings, we next sought a structural understanding of the ISRIB-stabilized human eIF2B decameric complex.

### ISRIB binds in a deep cleft, bridging the two-fold symmetric interface of the eIF2B decamer

We determined a near-atomic resolution structure of eIF2B bound to ISRIB by electron cryo-microscopy (cryoEM). We classified and refined a single consensus structure from 202,125 particles to an average resolution of 2.8 Å resolution, that varied from 2.7 Å in the stable core to >3.4 Å in the more flexible periphery (Fig. S3). The overall structure bears clear resemblance to the *S. pombe* two-fold symmetric decameric structure determined by X-ray crystallography (8). The symmetry interface comprises contacts between the α, β, and δ subunits, while the γ and ε subunits are attached at opposing ends (Fig. 2A-C). As in the *S. pombe* crystal structure, the catalytic HEAT domains of the ε subunits were not resolved, indicating their flexible attachment to the regulatory core. By contrast, densities for the “ear” domains of the γ subunits were resolved, but at a resolution that precluded atomic interpretation (Fig. 2B, Fig. S3).

**Fig. 2:**
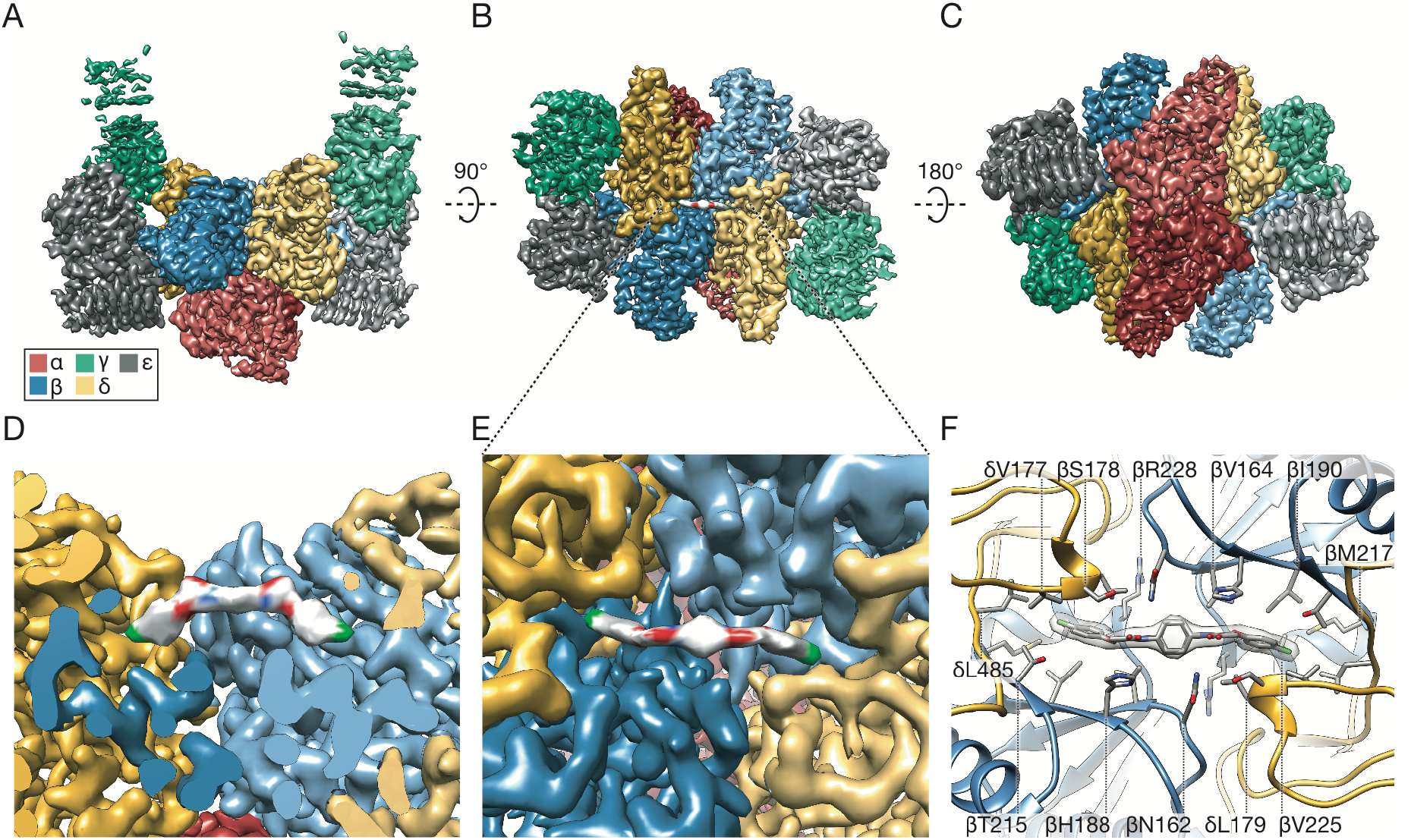
Near-atomic resolution reconstruction of ISRIB-bound eIF2B. (A-C) Three views of cryoEM density for eIF2B(αβγδε)_2_, colored in distinct shades for each subunit copy: red for α, blue for β, green for γ, gold for δ, and gray for ε (color code used throughout this manuscript). Density assigned to ISRIB depicted in CPK coloring: oxygens highlighted in red, nitrogens in blue and chlorines in green. The rotational relationships between the views depicted in A, B, and C are indicated. (D) Cross-section of (A), revealing view of the ISRIB binding pocket at the central decamer symmetry interface and density assigned to ISRIB CPK-colored by element. (E) Close-up view of density assigned to ISRIB and its binding pocket in (B) at the intersection of two β and two δ subunits. (F) Two conformers of ISRIB modeled into the density and all residues within a 3.7Å distance from the ligand rendered as sticks.

Importantly, we observed a clearly defined density consistent with the dimensions of ISRIB and not attributable to protein bridging the symmetry interface of the decamer (Fig. 2B, Fig. 2D-E, Fig. S4). Modeling suggests that ISRIB binds with its central cyclohexane ring in the expected low-energy chair conformation and with the side chains projecting to the same face of the cyclohexane ring and inserting the distal 4-chlorophenyl rings into deep binding pockets (Fig. 2D-F, Fig. S4). ISRIB’s "U-shaped" conformation may be stabilized by intramolecular N-H---O hydrogen bonding interactions between its amide nitrogen N-H bond and the aryl ether oxygens, possibly explaining why non-ether-linked congeners of ISRIB are much less potent (Fig. S5) (*28, 34*). The cryoEM density most likely corresponds with an average of at least two energetically equivalent ISRIB conformations related by 180° rotations about both N-C bonds to the cyclohexane ring (both depicted in Fig. 2F and Fig. S4–5). This superposition of two conformers accounts for the apparently symmetric density observed, even though in isolation each individual conformer is pseudo-symmetric (Fig. S4).

The N-terminal loop of the δ subunit contributes key residues to the binding pocket, and this loop differs significantly from the ligand-free *S. pombe* structure (*8*). Residues in this loop were previously shown to be important for ISRIB activity (*29*), including δV177 and δL179, which contribute directly to the hydrophobic surface of the binding pocket (Fig. 2F, Fig. S5). In addition, the δ subunits contribute δL485 to the hydrophobic wells that accommodate the halogenated benzene rings (Fig. 2F, Fig. S5). The center of the binding site comprises residues from the β subunit, including βN162 and βH188, which lie near ISRIB’s more polar functionality. In particular, one of the two C-H bonds at the glycolamide α-carbon is oriented perpendicular to the plane of the aromatic histidine ring (Fig. 2F, Fig. S5), suggesting a C-H-π interaction with βH188. Residues on the β subunits also make key contributions to the hydrophobicity of the deep wells, including βV164 and βI190.

Together these data suggest that ISRIB enhances incorporation of the α subunit into the decamer despite not making direct contacts with this subunit. Rather, ISRIB stabilizes the symmetry interface of the β-δ core, which in turn favors stable eIF2B(α_2_) binding. As such, ISRIB’s enhancement of GEF activity derives from its ability to promote higher-order holoenzyme assembly.

### Structural model predicts the activity of modified compounds and mutations

To validate the structural model, we synthesized ISRIB analogs bearing a methyl group at the a position of the glycolamide side chains. Two enantiomers, ISRIB-A19(*R,R*) and ISRIB-A19(*S,S*) were prepared (Fig. S6A) based on predicted steric clashes with residue δL179 for ISRIB-A19(*R,R*) or βH188 for ISRIB-A19(*S,S*) in the ISRIB binding pocket (Fig. 2F, Fig S5). As expected, neither enantiomer enhanced GEF activity *in vitro* or in cells (Fig. 3A, Fig. S6B), nor did they enhance the stability of purified decameric eIF2B (Fig. S6C). We next engineered eIF2B to accommodate the additional methyl groups on ISRIB-A19(*R,R*) by mutating δL179 to alanine. We tested the effects of both compounds on eIF2B(δL179A) by velocity sedimentation and GEF activity. As predicted, ISRIB-A19(*R,R*) stabilized formation of mutant decamers (Fig. 3B) and stimulated nucleotide exchange (Fig. 3C). Treatment with ISRIB-A19(*R,R*) activated eIF2B(δL179A) approximately three-fold (Fig. 3C, k_obs_ = 0.027 +/− 0.001 min^−1^), a similar fold-activation to eIF2B WT by ISRIB. By contrast and as predicted, ISRIB-A19(*S,S*) failed to activate eIF2B(δL179A) (Fig. 3C, k_obs_ = 0.007 +/− 0.001 min^−1^). Notably, in the absence of ISRIB analogs, eIF2B(δL179A) was five-fold less active than eIF2B (compare Fig. 3A and 3C, eIF2B k_obs_ = 0.04 +/− 0.009 min^−1^ and eIF2B(δL179A) k_obs_ = 0.008 +/− 0.002 min^−1^), identifying δL179A as a novel hypomorphic mutation and underscoring the importance of this surface for holoenzyme assembly.

**Fig. 3:**
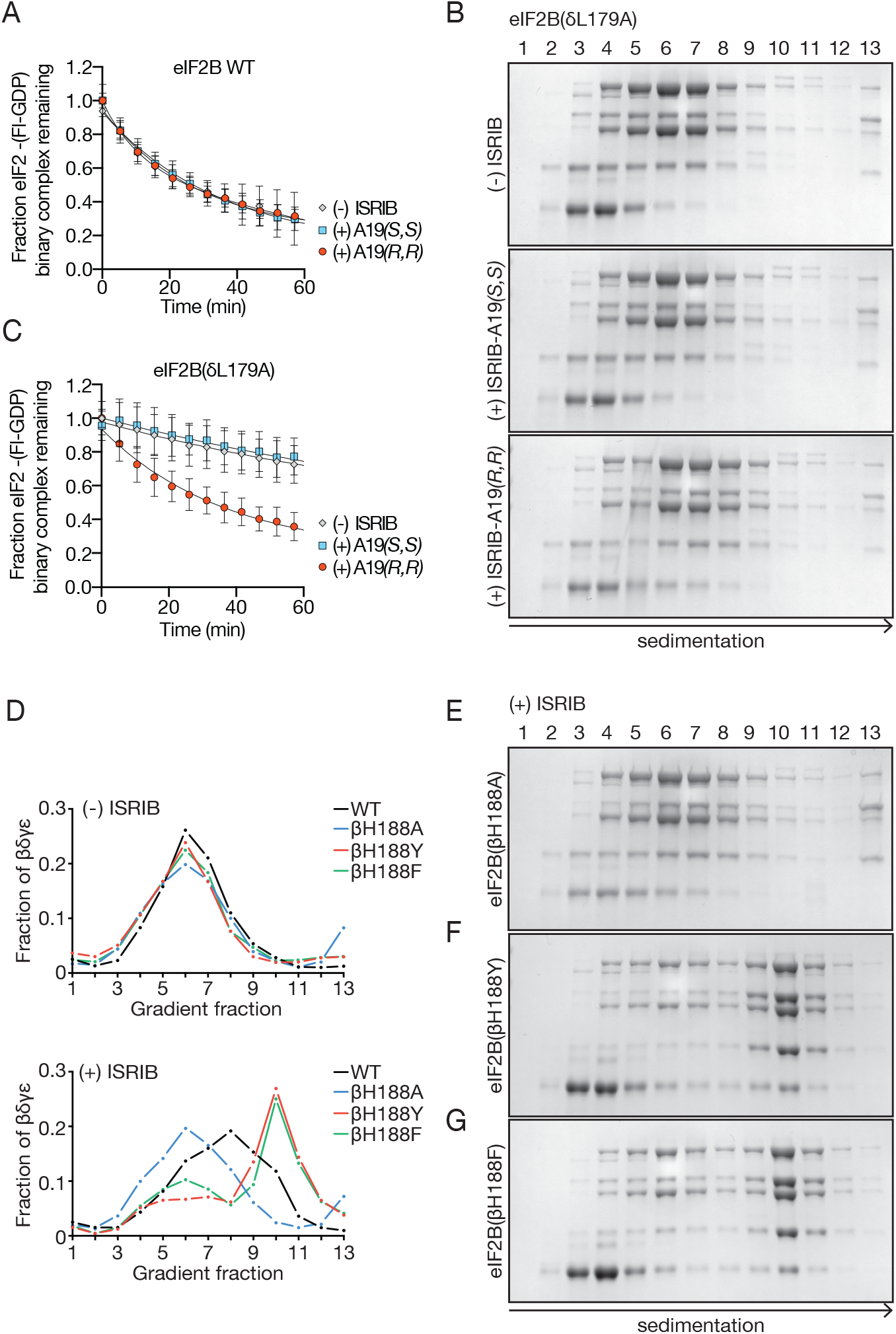
eIF2B structure predicts activity of ISRIB analogs. (A) GEF activity of assembled eIF2B(βγδε) and eIF2B(α_2_) in the presence and absence of ISRIB-A19(*R,R*) and ISRIB-A19(*S,S*). (B) Stability of decameric eIF2B(δL179A) in the absence of ISRIB (top), presence of ISRIB-A19(*S,S*) (middle), or presence of ISRIB-A19(*R,R*) (bottom) as assessed by velocity sedimentation on sucrose gradients. (C) eIF2B GEF activity of assembled eIF2B(βγδε) and eIF2B(α_2_) containing a δL179A mutation in the presence and absence of ISRIB-A19(*R,R*) and ISRIB-A19(*S,S*). (D) Quantification of eIF2B decamer stability gradients plotted as fraction of eIF2B(βγδε) present in each of lanes 1–13. eIF2B (for comparison from data shown in Fig. 1F), eIF2B(βH188A), eIF2B(βH188Y), eIF2B(βH188F) gradients are plotted in the presence (bottom panel) and absence (top panel) of 500 nM ISRIB. (E, F, G) Stability of decameric eIF2B(βH188A), eIF2B(βH188Y), and eIF2B(βH188F) in the presence of ISRIB as assessed by velocity sedimentation on sucrose gradients.

We next sought to verify the existence of a putative C-H-π interaction between βH188 and ISRIB by mutating βH188 to alanine. As predicted, ISRIB did not stabilize eIF2B(βH188A) decamers (Fig. 3D-E, Fig. S5). By contrast, mutating βH188 to an aromatic tyrosine or phenylalanine—which are predicted to sustain and likely enhance C-H-π interactions—did not impair ISRIB’s activity to stabilize decamers (Fig. 3D, Fig. 3F-G, Fig. S5). Rather, ISRIB stabilized eIF2B(βH188Y) and eIF2B(βH188F) decamers to an even greater extent than wild-type eIF2B decamers (Fig. 3D). Whereas ISRIB-stabilized wild-type eIF2B sedimented with a broad profile, indicating dissociation of the decamer through the course of sedimentation (Fig. 1F, Fig. 3D), ISRIB-stabilized eIF2B(βH188Y) and eIF2B(βH188F) formed a sharp symmetric peak in fraction 10, indicative of enhanced complex integrity through sedimentation, presumably due to enhanced C-H-π bonding interaction with ISRIB (Fig. 3D, Fig. 3F-G, Fig. S5).

### ISRIB induces dimerization of tetrameric eIF2B subcomplexes

Since ISRIB bridges the symmetry interface of the decamer without making direct contacts with eIF2B(α_2_), we sought to understand how the small molecule promotes eIF2B(α_2_) incorporation into the decamer. We imaged purified eIF2B(βγδε) tetramers in the presence and absence of ISRIB by cryoEM. In the presence of ISRIB, the images revealed a predominant species consistent with an octameric complex of eIF2B lacking the α subunits (Fig. 4A). By contrast, in the absence of ISRIB, the predominant species was consistent with a tetrameric complex divided along the symmetry axis of the octamer (Fig. 4B). In accordance with the ISRIB-dependent stabilization of the decamer by mutations in βH188 to other aromatic residues, βH188F and βH188Y mutants also stabilized the octamer in high salt conditions (Fig. S8). These images suggest a model in which ISRIB dimerizes eIF2B(βγδε) by “stapling” the tetramers together to form the octameric binding platform for a subunit binding, consistent with the architecture of the ISRIB-bound decamer.

**Fig. 4:**
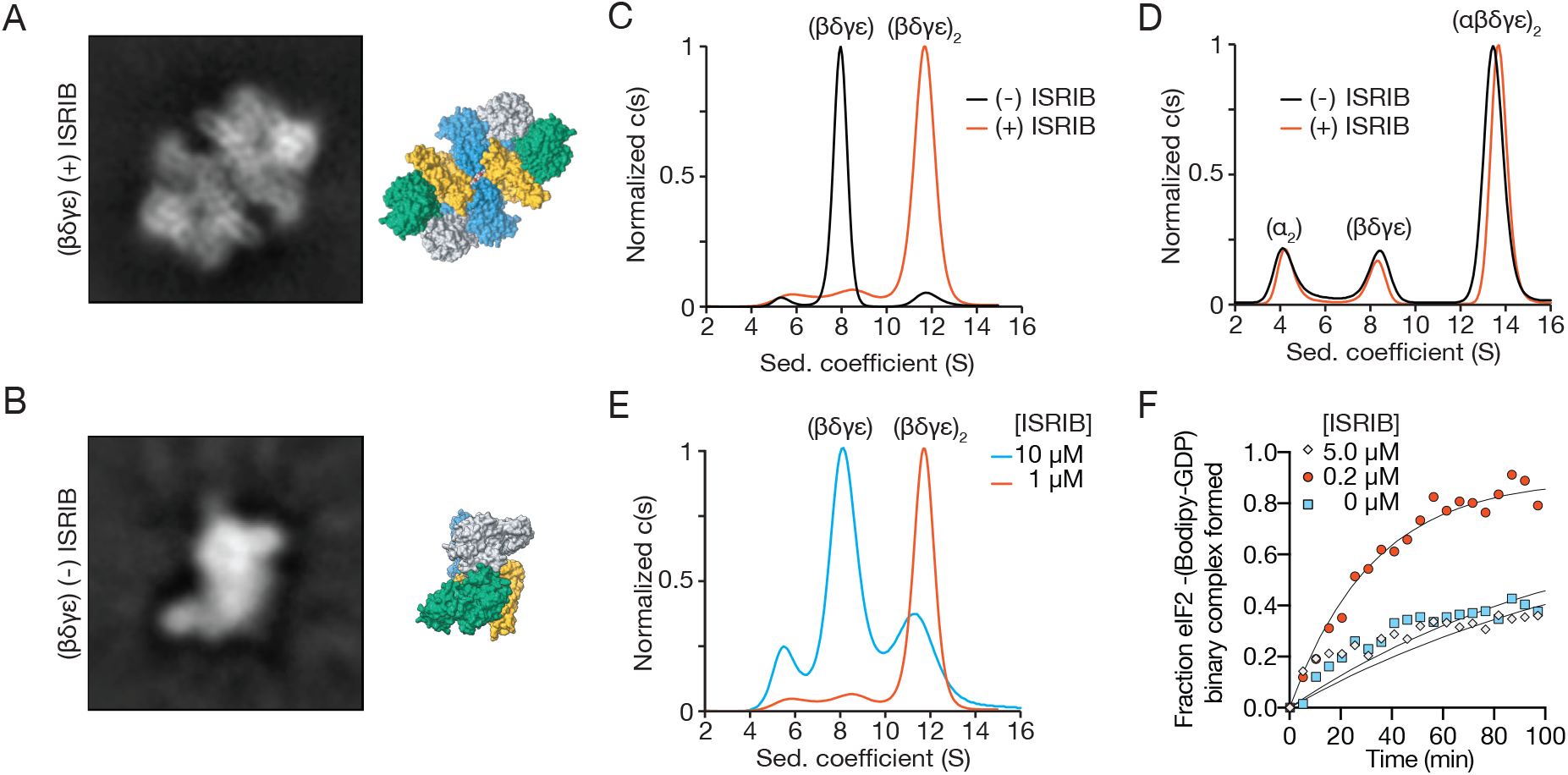
ISRIB induces dimerization of tetrameric eIF2B subcomplexes. The most abundant 2D class averages from cryoEM imaging of eIF2B(βγδε) in the presence (A) and absence (B) of ISRIB. (C) Characterization of eIF2B(βγδε) by sedimentation velocity analytical ultracentrifugation. eIF2B(βγδε) (1 μM) was analyzed in the presence and absence of 1 μM ISRIB. (D) Mixture of 1 μM eIF2B(βγδε) and 500 nM eIF2B(α_2_) characterized by analytical ultracentrifugation in the presence and absence of 1 μM ISRIB. (E) eIF2B(βγδε) (1 μM) characterized by analytical ultracentrifugation in the presence of 1 μM or 10 μM ISRIB. (F) GEF activity of eIF2B(βγδε), here at a higher 100nM concentration to facilitate comparison of 0, 0.2, and 5 μM ISRIB.

We next substantiated eIF2B(βγδε) dimerization by analytical ultracentrifugation under physiological salt conditions. In the absence of ISRIB, eIF2B(βγδε) sedimented as a predominant 8.0S peak and a minor 11.7S peak, corresponding to eIF2B(βγδε) and eIF2B(βγδε)_2_, respectively (Fig. 4C). By contrast, in the presence of ISRIB, we observed a dramatic increase in the 11.7S peak, demonstrating ISRIB’s role in stabilizing the eIF2B(βγδε)_2_ octamer. Together with the observation that eIF2B(βγδε) has greater activity in the presence of ISRIB (Fig. 1I), these data show the importance of octamer assembly in activating GEF activity.

Dimerization of eIF2B(βγδε) effectively doubles the surface area for eIF2B(α_2_) binding, suggesting that the ISRIB-enhanced incorporation of eIF2B(α_2_) into the decamer originates from ISRIB’s ability to shift the tetramer/octamer equilibrium. To test this prediction, we combined eIF2B(α_2_) and eIF2B(βγδε) in the presence and absence of ISRIB and assessed decamer assembly by analytical ultracentrifugation. Under the high protein concentrations used in these assays, we observed a predominant peak corresponding to the assembled eIF2B decamer at 13.6S both in the presence and absence of ISRIB, together with minor peaks corresponding to unincorporated eIF2B(βγδε) at 8.0S and eIF2B(α_2_) at 4.1S (Fig. 4D). Importantly, we did not observe an octamer peak, suggesting the octamer has a high affinity for eIF2B(α_2_) and assembles the fully assembled decamer under these conditions. Together with the cryoEM images, these data demonstrate that eIF2B(α_2_) and ISRIB synergistically promote dimerization of eIF2B(βγδε).

Given that ISRIB binds across the eIF2B(βγδε)_2_ interface such that each tetramer contributes half of the ISRIB binding site, we reasoned that high ISRIB concentrations may occupy half-sites within the tetramers and interfere with octamer formation. Indeed, ISRIB promoted eIF2B(βγδε)_2_ assembly at 1 μM but failed to do so at 10 μM (Fig. 4E). Similarly, ISRIB stimulated GEF activity of eIF2B(βγδε) at 200 nM but failed to do so at 5 μM (Fig. 4F). Importantly, the high ISRIB concentrations used in this assay did not reduce GEF activity below that of eIF2B(βγδε), demonstrating that the effect did not result from non-specific enzymatic inhibition.

### Loss and gain-of-function dimerization mutants resist or bypass the effects of ISRIB

To visualize the determinants of octamerization, we highlighted the solvent-excluded surface area along the symmetry interface of the β and δ subunits in adjacent tetramers (Fig. 5A-B, light yellow, light blue, green) and labeled the residues of the ISRIB binding pocket on this surface (Fig. 5A-B, gray). The tetramer-tetramer contact residues form a thin strip along each neighboring β and δ subunit. Most of the β subunit residues contact the δ subunit across the symmetry interface, while a small number of residues also cement β-β’ contacts. Of these, βH160 and βR228 reside at the junction of β-β’ and β-δ’ subunits, suggesting that they play key roles in stabilizing the octamer. Accordingly, we observed that mutation of βH160 to aspartic acid, which we predicted would be repulsed by δD450, completely precluded octamer assembly. Analytical ultracentrifugation of eIF2B(βγδε) containing the βH160D mutation revealed a sharp tetramer peak at 7S both in the absence and presence of ISRIB (Fig. 5C), and ISRIB was unable to enhance GEF activity for this mutant (Fig. 5D). These observations indicate that the effect of this mutation on octamerization cannot be overcome by ISRIB binding, despite the fact that ISRIB binding buries an additional ~11% of solvent-exposed surface area—an increase from 3420 Å^2^ to 3790 Å^2^ —upon stapling of tetramers (Fig. 5A-B).

**Fig. 5:**
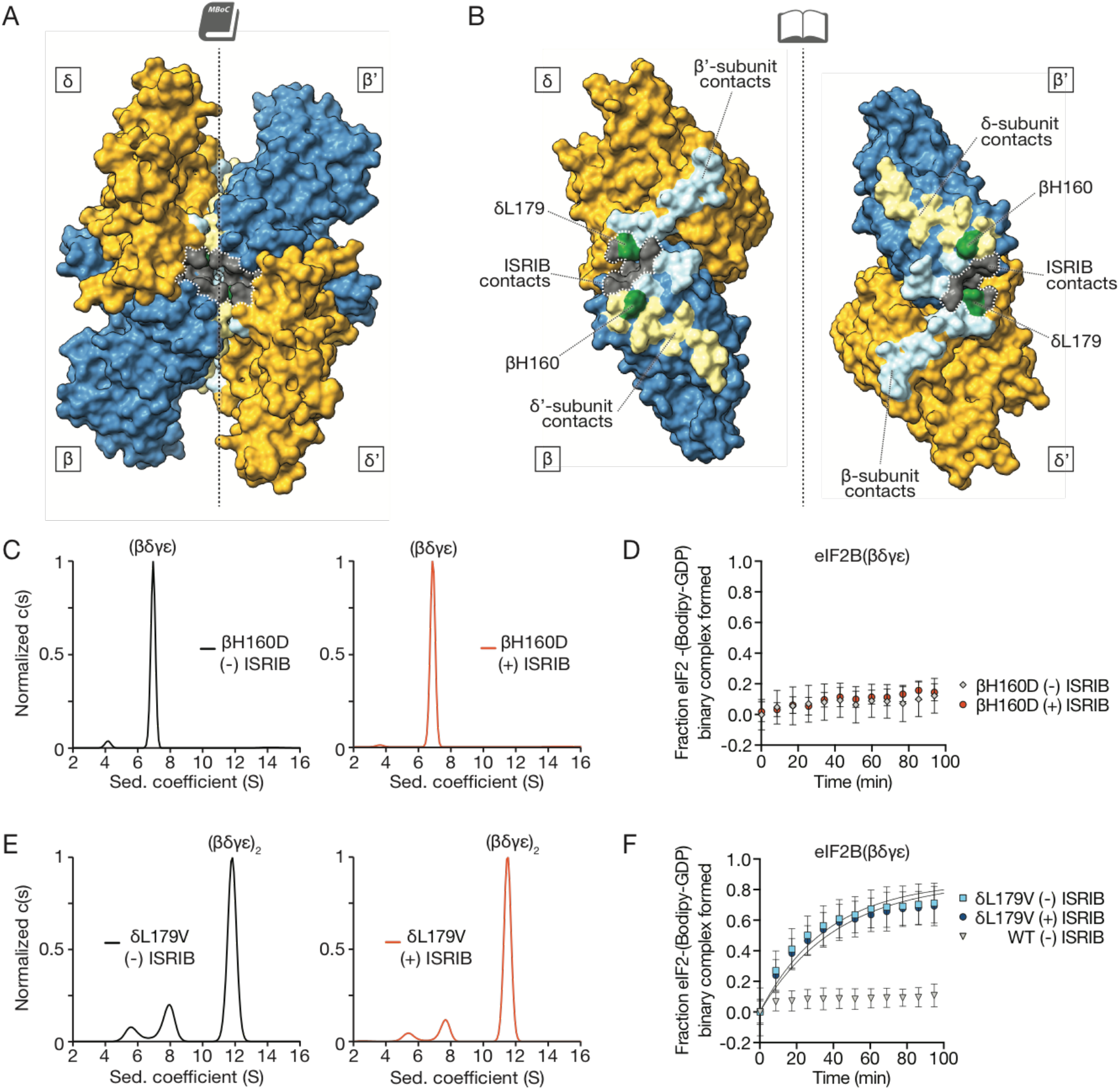
Loss- and gain-of-function dimerization mutants resist or bypass the effects of ISRIB. (A) Surface rendering of core eIF2Bβ (blue) and eIF2Bδ (gold) subunits with residues contacting ISRIB highlighted in gray and with dimer interface indicated by dashed line. Interface residues are highlighted in a lighter hue of the colors of the contacting subunits. (B) Open-book view of the dimer-dimer interface, such that each β and δ subunit is rotated by 90°. βH160, in green, contacts both β’ and δ’; δL179, also in green, contacts both β’ and ISRIB. (C) Characterization of 1 μM eIF2B(βγδε) containing a βH160D mutation in the presence (right) and absence (left) of 1 μM ISRIB by analytical ultracentrifugation. (D) GEF activity of eIF2B(βγδε) containing a βH160D mutation in the presence and absence of ISRIB. (E) Characterization of 1 μM eIF2B(βγδε) containing a δL179V mutation in the presence (right) and absence (left) 1 μM ISRIB by analytical ultracentrifugation. (F) GEF activity of eIF2B(βγδε) containing a δL179V mutation in the presence and absence of ISRIB.

Serendipitously, we also identified a gain-of-function mutation in eIF2B. We initially engineered a δL179V mutation alongside the δL179A mutation used above to accommodate the methylated analog ISRIB-A19(*R,R*) (Fig. 2F, Fig. S5). To our surprise, we discovered that the predominant species of δL179V-eIF2B(βγδε) sediments as a remarkably stable octamer in the absence of ISRIB (Fig. 5E). GEF activity assays revealed that δL179V-eIF2B(βγδε)_2_ was five-fold more active than the wild-type octamers formed in the presence of ISRIB, and was not further activated by ISRIB (compare Fig. 5F and Fig. 1I, eIF2B(δL179V) k_obs_ = 0.027 +/− 0.001 min^−1^, eIF2B(δL179V) + ISRIB k_obs_ = 0.024 +/− 0.001 min^−1^, WT + ISRIB k_obs_ = 0.005 +/− 0.001 min^−1^). Together with the ISRIB-bound structure, these mutants indicate that the major contribution of ISRIB to increased GEF activity lies at the step of tetramer dimerization and assembly of the bipartite surface for a subunit homodimer binding (Fig. 6).

**Fig. 6:**
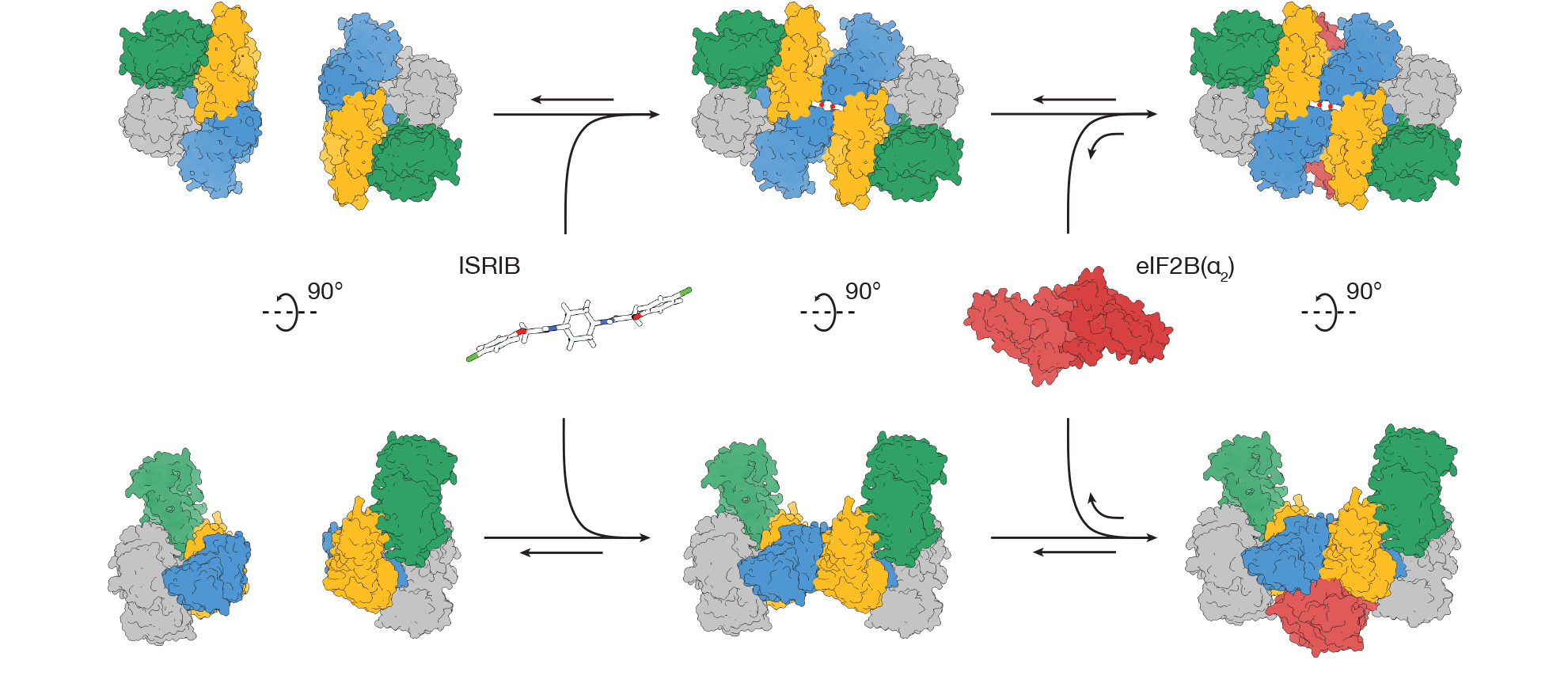
Model for ISRIB’s mechanism of action. ISRIB staples together tetrameric eIF2B(βγδε) subcomplexes, building a more active eIF2B(βγδε)_2_ octamer. In turn, the ISRIB-stabilized octamer binds eIF2B(α_2_) with greater affinity, enhancing the formation of a fully-active, decameric holoenzyme.

## Discussion

We determined the first structure of human eIF2B at sufficiently high resolution to characterize the binding-site and coordination of a new-class of small molecule with therapeutic potential. The atomic model of ISRIB-bound eIF2B reconciles structure-activity-relationships described previously (*28, 34*), proved predictive of both loss- and gain-of-function mutations, and greatly enables the rational design of new small molecule modulators of eIF2B activity. The structure provides an intuitive view of how ISRIB activates nucleotide exchange: ISRIB stabilizes the active decameric form of the eIF2B holoenzyme by stapling the constituents together across a 2-fold symmetry axis.

Given that a catalytic residue essential for nucleotide exchange resides in the still unresolved HEAT repeat of the ε subunit, how does assembly of the decameric holoenzyme enhance activity? Crosslinking studies suggest that eIF2 binds across the decameric interface, engaging the eIF2B α subunit, and β and δ subunits from opposing tetramers (*8*). It is therefore reasonable to surmise that decamer assembly creates a composite surface for eIF2 binding that allows the flexibly attached HEAT domain to reach and engage its target. While we consider it likely that the effects of ISRIB binding can be explained by the degree of holoenzyme assembly, additional ligand-induced allosteric changes may also contribute to its activity.

These observations provide a plausible model for ISRIB’s ability to ameliorate the inhibitory effects of eIF2α phosphorylation on ternary complex formation. Using purified components we show both mechanistically and structurally how ISRIB staples tetrameric building blocks together into an octamer, which enhances activity three-fold, and forms a platform for association of the dimeric a subunits. The integrated effect of these sequential steps is an order of magnitude enhancement of activity. The inhibition resulting from a limiting amount of phosphorylated eIF2 would be reduced by the surplus of GEF activity provided by ISRIB. By contrast, we noted that an excess of ISRIB poisons the assembly reaction by saturating half-binding sites on unassembled tetramers. Together, these observations indicate that, within its effective concentration range, ISRIB will enhance ternary complex formation even in unstressed conditions, opening an untapped reservoir of additional enzymatic capacity. We surmise that *in vivo* these activities are realized near the equilibrium points of the assembly reactions for the holoenzyme, allowing for ISRIB’s observed phenotypic effects. Thus, eIF2B is poised to integrate diverse signals that impact translation initiation. Phosphorylation of eIF2 may be just one of many mechanisms for modulating its activity. Post-translational modifications, expression of other modulatory components, or binding of yet-to-be-identified endogenous ligands to the ISRIB binding pocket or elsewhere are likely to modulate eIF2B activity under varying physiological conditions. Understanding the different modes of regulation of this vital translational control point will be of particular importance in the nervous system where ISRIB was shown to have a range of impressive effects.

## Acknowledgments

We thank Graham Pavitt for the GP6452 yeast strain used in the purification of eIF2. We thank Jirka Peschek, Elif Karagöz, Robert Stroud, James Fraser, Geeta Narlikar, Ron Vale, Axel Brilot, Nicole Schirle Oakdale, Nathaniel Talledge, Pearl Tsai, Ni Mu, Joseph Choe, Christopher Upjohn, and the Walter and Frost labs for reagents, technical advice and helpful discussions. We thank Michael Braunfeld, David Bulkley, and Alexander Myasnikov of the UCSF Center for Advanced CryoEM and Daniel Toso and Paul Tobias of the Berkeley Bay Area CryoEM Facility, which are supported by in part from NIH grants S10OD020054 and 1S10OD021741 and the Howard Hughes Medical Institute (HHMI). We also thank Zhiheng Yu, Rick Huang, and Chuan Hong of the CryoEM Facility at the Janelia Research Campus of the HHMI. We thank the QB3 shared cluster and NIH grant 1S10OD021596-01 for computational support. This work was supported by funding to AF from a Faculty Scholar grant from the HHMI, the Searle Scholars Program, and NIH grant 1DP2GM110772-01, and by funding to PW from Calico Life Sciences LLC, the Rogers Family Foundation, the Weill Foundation, and the HHMI. AF is a Chan Zuckerberg Biohub Investigator, and PW is an Investigator of the HHMI. PW and ARR are listed as inventors on a patent application describing ISRIB and analogs. Rights to the invention have been licensed by UCSF to Calico.

## Supplementary Materials

Materials and Methods

Figures S1-S8

Tables S1-S3

